# Strategic decision support for long-term conservation management planning

**DOI:** 10.1101/2021.06.17.448842

**Authors:** Eric S. Abelson, Keith M. Reynolds, Patricia Manley, Steven Paplanus

## Abstract

Forward thinking conservation-planning can benefit from modeling future landscapes that result from multiple alternative management scenarios. However, long-term landscape modeling and downstream analyses of modeling results can lead to massive amounts of data that are difficult to assemble, analyze, and to report findings in a way that is easily accessible to decision makers. In this study, we developed a decision support process to evaluate modeled forest conditions resulting from five management scenarios, modeled across 100 years in California’s Lake Tahoe basin; to this end we drew upon a large and complex hierarchical dataset intended to evaluate landscape resilience. Trajectories of landscape characteristics used to inform an analysis of landscape resilience in the Lake Tahoe basin were modeled with the spatially explicit LANDIS-II vegetation simulator. Downstream modeling outputs of additional landscape characteristics were derived from the LANDIS-II outputs (e.g., wildlife conditions, water quality, effects of fire). The later modeling processes resulted in the generation of massive data sets with high dimensionality of landscape characteristics at both high spatial and temporal resolution. Ultimately, our analysis distilled hundreds of data inputs into trajectories of the performance of the five management scenarios over the 100-year time horizon of the modeling. We then evaluated each management scenario based on inter-year variability, and absolute and relative performance. We found that the management scenario that relied on prescribed fire, outperformed the other four management approaches. Both these results, and the process that led to them, provided decision makers with easy-to-understand results based on a rational, transparent, and repeatable decision support process.

**One sentence description:** We present a novel approach to employ decision support tools for conservation over long time horizons.

## 1. Introduction

Conservation and restoration planning across forested landscapes has become exponentially more complex and challenging over the past decade, primarily because of the effects of human-based landscape alteration and impacts of climate change across the full spectrum of ecological and social systems (e.g. (Noss, 2001)). Forested ecosystems provide essential services and benefits to society, and concomitantly those services and benefits require substantial management inputs to be achieved and maintained (Sutherland et al., 2014). Resilience has become a beacon for restoration and conservation outcomes, which reflects the desire to maintain characteristic composition, processes, and functions of ecosystems over time and in response to perturbations (Walker et al., 2004). Understanding future trends in climate, and their potential impacts on forest ecosystems, is now necessary to inform even near-term project planning, and increasing the pace and scale of restoration to improve climate readiness and ecological resilience is a widely shared objective (Belote et al., 2018; Lawler et al., 2015). As management planning projects increase in spatial scale and extent, so do the potential impacts and potential benefits on a wide range of resource and societal values (Messier et al., 2015). Hence, management projects are increasingly grappling with large spatial scales, multiple natural resource objectives, a spectrum of societal values, and multi-decadal temporal dynamics with an uncertain climate future.

The Lake Tahoe basin is a prime example of a landscape where high value ecological and social outcomes hang in the balance as climate changes. Recent climate assessments for California (Bedsworth et al., 2018) and the Lake Tahoe basin (California Tahoe Conservancy, n.d.) have raised substantial concerns for the health of its forests, conservation of biodiversity, and the future of the renowned clarity of Lake Tahoe. Agencies, academic institutions, and non-governmental organizations came together to form the Lake Tahoe West Restoration Partnership to gain a better understanding of conservation options and expedite management actions to improve climate readiness of forest ecosystems on the west side of the Lake Tahoe basin. A science team was convened that included experts in the fields of forest management, ecology, hydrology, wildlife biology, and fire science to conduct an integrated modeling approach to understanding ecological and social outcomes associated with multiple long-range management scenarios.

The ability to evaluate multiple resource and societal outcomes across unique future management scenarios is becoming an essential capacity to inform and support project design and planning decisions. However, decision support tools that can incorporate large spatial scales, long time frames, multiple management scenarios, and composite suites of resource and societal values are limited. The Ecosystem Management Decision Support System (EMDS) (Reynolds and Hessburg, 2014) is one of the few tools that has demonstrated capability in handling these types of multi-dimensional land management applications (Reynolds et al., 2014). Past applications of EMDS have predominantly used current landscape condition, desired landscape condition, and the societal/ecological costs and benefits to identify areas within landscapes that, if treated, would improve overall conditions. From its earliest design specifications up to the present time, EMDS is best known as a *spatial* decision support system (SDSS) for environmental analysis and planning (Reynolds and Hessburg, 2014). EMDS applications addressing these challenges assess large amounts of data (that have a spatial component), ultimately resulting in tangible guidance for decision makers (Cleland et al., 2017; Hessburg et al., 2013; Reynolds, 2001; Reynolds et al., 2009; Walker et al., 2007). However, EMDS is a highly flexible tool that can be used to account for future trends and alternative management scenarios (whether desired output is spatial or aspatial).

In this paper, we present a novel application of the EMDS system (Reynolds and Hessburg, 2014) to support long-term strategic planning for ecosystem resilience in the Lake Tahoe West (LTW) region of the Lake Tahoe basin. The LTW application of EMDS is novel in four main ways. First, evaluation of ecosystem resilience in LTW is aspatial: while our logic-based evaluation of ecosystem resilience is typical of spatial EMDS applications (insofar as the logic engine is used to interpret and synthesize numerous complex, abstract, and high-dimensional ecosystem attributes representing facets of ecosystem resilience), ultimate results are aspatial metrics of management scenario performance over time. Second, LTW results consider a long time-horizon (i.e., a century): while EMDS has traditionally examined existing on-the-ground conditions at a single point in time, we have used EMDS in this application to prospectively evaluate forest conditions over a century. Third, multiple modeled future conditions are assembled into a single analysis: in the LTW project, EMDS contrasts future forest conditions for five unique landscapes, based on five distinct forest management approaches, over a 100-year time span. Finally, the LTW application of EMDS is also novel with its demonstration of a new automated workflow feature implemented in EMDS, which involves iterating logic model and decision model processing over multiple alternative management scenarios and time steps.

The present study demonstrates how a diverse array of ecological conditions in an ecosystem can be assessed for ecological resilience over time with tractable output for decision-makers; this is an analytical problem with high dimensionality in space, time, and environmental complexity that, as we shall demonstrate, can be distilled down, through a series of analytical steps, to a relatively simple graphical representation in which trajectories of resilience performance for multiple strategic management scenarios for improving ecosystem resilience are compared and contrasted over the next century. A full description of the complete set of models needed to assess ecosystem resilience in LTW is impractical to present in a single paper, so the complete resilience analysis for LTW is presented in two reports. In this first paper, we focus specifically on analysis of resilience of ecological conditions to demonstrate the details of the methodology. The second paper (Abelson et al., n.d.) builds on the present work to provide a broader interpretation of the performance of ecosystem resilience by also accounting for the social and economic dimensions of ecosystem resilience.

## 2. Materials and methods

### 2.1. Study area

The west side of the Lake Tahoe Basin, California served as the study area for these analyses. The study area consists of 23,882 ha between Emerald Bay to the south and Squaw Valley to the north (Figure 1). All lands were included in the analysis, but management scenarios (i.e., fuel reduction activities) pertained to public lands only (National Forest System lands of US Department of Agriculture Forest Service and California State lands).

**Figure 1:**
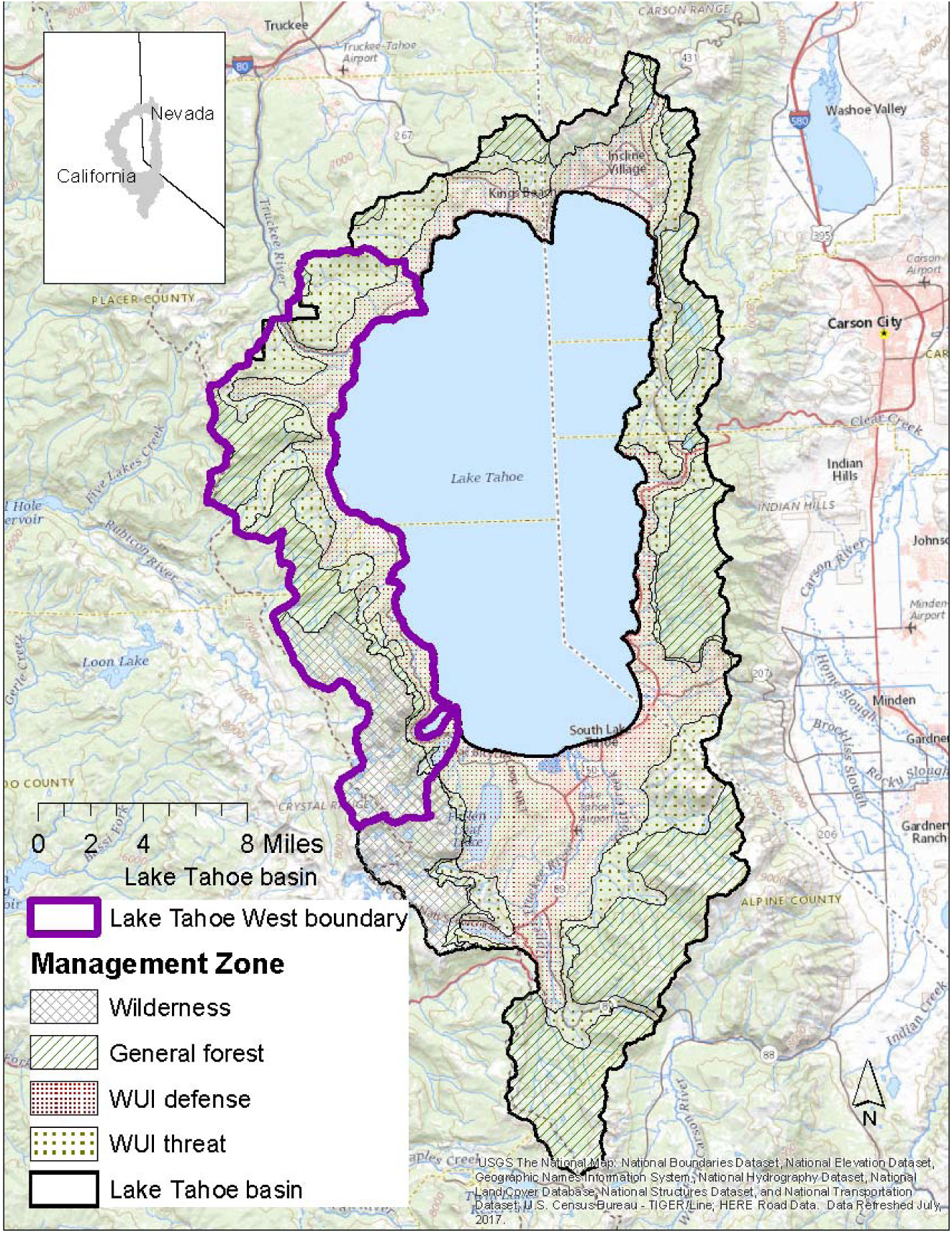
Study area is the western section (outlined in purple) of the Lake Tahoe basin (outlined in black).

The Lake Tahoe basin was selected as focus of this application because it is a high value, at-risk landscape, and it has stakeholder and tribal communities that are highly engaged in informing management toward restoring ecological resilience. Decision makers need to not only understand the perspectives of a multitude of stakeholders and partners, but to the degree possible they need to understand the complexities of a myriad of interacting factors that affect outcomes, including ecological and social values that hang in the balance. The west side of Lake Tahoe is particularly vulnerable to ecological impacts from fire and climate change because it is has the most biodiverse and productive forests, so they are dense, and it has the highest wildland-urban interface, so the majority (>60%) of the west side of the basin is WUI and the threat of fire is high. In other words, it is both a literal and figurative tinder box – just the sort of circumstance that decision support tools are designed to address.

### 2.2. Overview of the decision support process for assessing ecological resilience

The LTW decision support process was accomplished in five main steps: Step 1 was to establish goals, including our spatial and temporal scope (Section 2.3). Step 2 was focused on management strategies, ecological topics of interest, and data needs (Section 2.4) while Step 3 focused on ecosystem modeling and data acquisition (Section 2.5). Step 4 was the heart of our decision support process and consisted of summarizing detailed data inputs into general topic areas using a NetWeaver logic model (Section 2.6) and then assessing all topics using a multi-criteria decision model (MCDM; Section 2.7) to prioritize and rank management strategies against each other. Finally, in Step 5, we present both results directly from the decision model in conjunction with summary statistics and visualizations (Section 2.8).

The logic models (Section 2.6) were used to interpret and synthesize numerous data inputs into higher order logic topics. Logic outputs were then entered into a decision model that was used to examine all available evidence, using the lens of a decision maker, to provide tractable guidance for understanding the role that management plays in promoting resilient ecological conditions in LTW. To facilitate running the several logic models and the decision model at each of 10 time steps, we implemented a workflow in the new EMDS workflow editor to automate the processing (Supplements 1 and 2).

### 2.3. Project goals

To establish project goals and desired output, a group of core LTW stakeholders was convened that was open to any organization with interest in the basin, and included representatives from primary federal and state agencies operating in the west side of the basin (US Department of Agriculture, Forest Service (USFS), Lake Tahoe Basin Management Unit, Tahoe Regional Planning Agency, California State Parks, Lahonton Water Quality Control Board, California Tahoe Conservancy, and CalFire. Nearly 20 other interest groups representing a wide array of organizations and institutions were also represented in the Lake Tahoe West stakeholder group. The objective of this project was to help decision makers evaluate management strategies for biomass removal and fire suppression in terms of each strategy’s ability to promote forest environmental quality and thereby ecological resilience (hereafter, for brevity, we refer to environmental quality). Forest conditions are dynamic and fluctuate over time, so, to better understand how divergent management approaches and climate change are likely to influence forest conditions over time, we modeled 100 years of prospective conditions in the Lake Tahoe Basin for each management strategy. We hypothesized that alternative fire suppression policies and amounts and types of biomass reduction (i.e., fire-based versus non-fire-based biomass removal) would differentially influence environmental quality and ecological resilience over time.

### 2.4. Management strategies, ecological topic areas, and data needs

Five management scenarios were designed and evaluated for their performance with respect to achieving environmental quality across the LTW landscape. Each management scenario was designed to represent a specific management perspective and to highlight the possible differences in environmental quality over a 100-year timespan. The five modeled management scenarios selected by LTW stakeholders were:

**Scenario 1. Fire suppression only** – No management other than suppressing natural or arson-caused fire ignitions.

**Scenario 2. Wildland-urban interface focus** - Forest thinning is restricted to the wildland-urban interface (WUI, areas near human habitation) to provide defensible space around structures and private property. Activities in scenario 2 largely reflect current management practices in the Lake Tahoe basin.

**Scenario 3. Forest thinning-based approach** – This scenario expands forest thinning in the WUI to areas outside the WUI using mechanical biomass removal methods.

**Scenario 4. Fire-based approach** – This scenario expands forest thinning in the WUI to areas outside the WUI using prescribed fire for one month per year.

**Scenario 5. Intensive fire-based approach** - This scenario expands forest thinning in the WUI to areas outside the WUI with a more intense use of, compared to scenario 4, prescribed fire (i.e., through the entire year).

To evaluate environmental quality under each of the five management scenarios, the two senior authors facilitated the design of the required logic models and decision model, working with the LTW stakeholders. Further details about data sources and model designs based on stakeholder input are presented in Sections 2.5, 2.6, and 2.7.

### 2.5. Landscape modeling and data inputs to the logic and decision models

Research by other members of the LTW science team modeled landscape characteristics (e.g., vegetation, fire, and beetle dynamics) annually for 100 years using LANDIS-II (*LANDIS-II*, n.d.; Scheller et al., 2007). Each of the five scenarios were modeled separately, producing a representation of the LTW landscape for each year and scenario combination (i.e., 500 modeled landscapes). All data used in our analysis were derived from LANDIS-II simulations (Supplement 3). The LANDIS-II forest landscape dynamics model projected spatial attributes of trees and shrubs at annual time steps and at 1-ha spatial resolution, considering prior-year forest conditions (or current conditions in year one), a changing climate, management activities, and stochastic events such as fire. Vegetation properties were simulated for a 100-yr period (from 2010 to 2109) for each of the five management scenarios described in Section 2.4. LANDIS-II modeling accounted for climate based on changes predicted by a representative concentration pathway of 4.5 and output from four global circulation models identified as the most likely in the most recent, California’s Fourth Climate Change Assessment (Bedsworth et al., 2018). LANDIS-II simulation outputs then served as input data for follow-on (i.e., secondary) modeling by teams of LTW scientists addressing landscape resilience characteristics for wildlife biology, hydrology, fire ecology, landscape ecology (e.g., vegetation), economics, and forestry (Long et al., 2020). The latter secondary outputs were similarly calculated at annual time steps and 1-ha spatial resolution.

Although EMDS is designed as a spatial decision support framework and LANDIS-II output were spatially explicit, the LTW project goals were aspatial in nature. Spatial attributes of modeled landscape characteristics were summarized across the spatial extent of the study area, thus removing the spatial dimension (Long et al., 2020) before data were input into the logic (Section 2.6) and decision (Section 2.7) models.

### 2.6. Logic models

We used logic-based processing (Guarino, 1998) to reduce the dimensionality of landscape characteristics to a set of four high-order landscape characteristics of most interest to land managers (Abelson, 2021; Supplements 1 and 4). The logic models in the LTW project assembled and assessed 33 complex environmental data inputs into four higher order, more abstract logic topics, which then served as decision model input (Section 2.7). The four logic models addressing aspects of ecological resilience of environmental quality were: wildlife conservation, quality water, upland vegetation health, and functional fire. Each logic model provided an interpretation of raw data; for example, soil aerator functional redundancy, early-seral beta diversity, and 11 other sub-topics were evaluated by a logic model into a single metric representing wildlife conservation for each time point and management scenario. Descriptions of the individual logic models are presented in Section 2.6.1.

The logic modeling component in EMDS, NetWeaver Developer from Rules of Thumb, Inc. (*NetWeaver*, 2020), evaluates data against a logic model that provides a formal specification for interpreting data and synthesizing information (Saunders and Miller, 2014). The logic processor readily supports design of logic specifications for the types of large, complex, and abstract problems typically posed by contemporary environmental management issues, such as ecosystem integrity (Cleland et al., 2017) or ecosystem resilience. The LTW project used logic models as pre-processors of complex problem types to distill down high dimensional or highly nonlinear information for improved use in subsequent decision models (Section 2.7)). The semantics and syntax of logic-based reasoning provided a modeling environment in which our large, interdisciplinary teams of scientists and managers could collaboratively assemble their respective facets of knowledge about systems into a relatively holistic representation of complex phenomena such as ecosystem integrity or resilience (Reynolds and Hessburg, 2014).

In the LTW project, we ran each of the four NetWeaver logic models 50 times; one model run for each scenario and decade combination (five scenarios by 10 time points). This produced 200 NetWeaver outputs; one output value for each of the four models at each of the 10 time-points and five management scenarios. NetWeaver output values vary between -1 and 1. A NetWeaver value of 1 reflects ecological conditions that are fully consistent with optimal resilience for the LTW forest landscape. A NetWeaver value of -1 reflects ecological conditions that are antithetical to resilience. To improve ease of interpreting NetWeaver values and Criterion DecisionPlus decision model values in concert, NetWeaver values were rescaled to the range of [0, 1] so that they had the same range as decision model results.

#### 2.6.1. Logic model structure & data description

NetWeaver logic models have a network architecture in which each NetWeaver topic is dependent on subtopics and subtopics, in turn, may also have subtopics. The four primary LTW project logic models for inferring ecological resilience of environmental quality, and their subtopics, are described in the following four subsections (2.6.1.1 to 2.6.1.4) and can be seen in complete detail in the NetWeaver model HTML documentation (Abelson, 2021; full NetWeaver model is available in Supplement 4). Throughout the following four subsections, we use the term “evaluate” in the specific context of data (as opposed to evaluating logic topics) as a shorthand to indicate evaluation of data by a fuzzy membership function (FMF, Miller and Saunders 2002, detailed description is available in Supplement 1 and values used in models presented here are available in Supplement 5). Ultimately, output from NetWeaver consisted of four values representing conditions for each of the four logic model topics by five scenarios and 10 decades (i.e., 200 values). These valuations were then used as input for the decision model.

##### 2.6.1.1. Wildlife conservation

The wildlife conservation topic consists of two levels of subtopics (Figure 2). UNION operators are used exclusively within the wildlife conservation topic. In the LTW project, wildlife conservation conditions are evaluated based on four subtopics, including species richness, ecological function, species diversity, and apex predators. Species richness evaluates the proportion of species for which greater than 70% of their 2010 habitat area is maintained. Ecological function is based on six subtopics; each subtopic evaluates the proportion of species (in the given category) in which 70% of their 2010 habitat area is maintained. Species diversity is based on three subtopics, with each evaluating the proportion of species that have habitat in early-, mid- and late-seral forest at any given time step and the total number of possible species. The Apex predator subtopic evaluates the proportion of predicted number of territories, by species, to the total possible number of territories that could be supported by the landscape.

**Figure 2.**
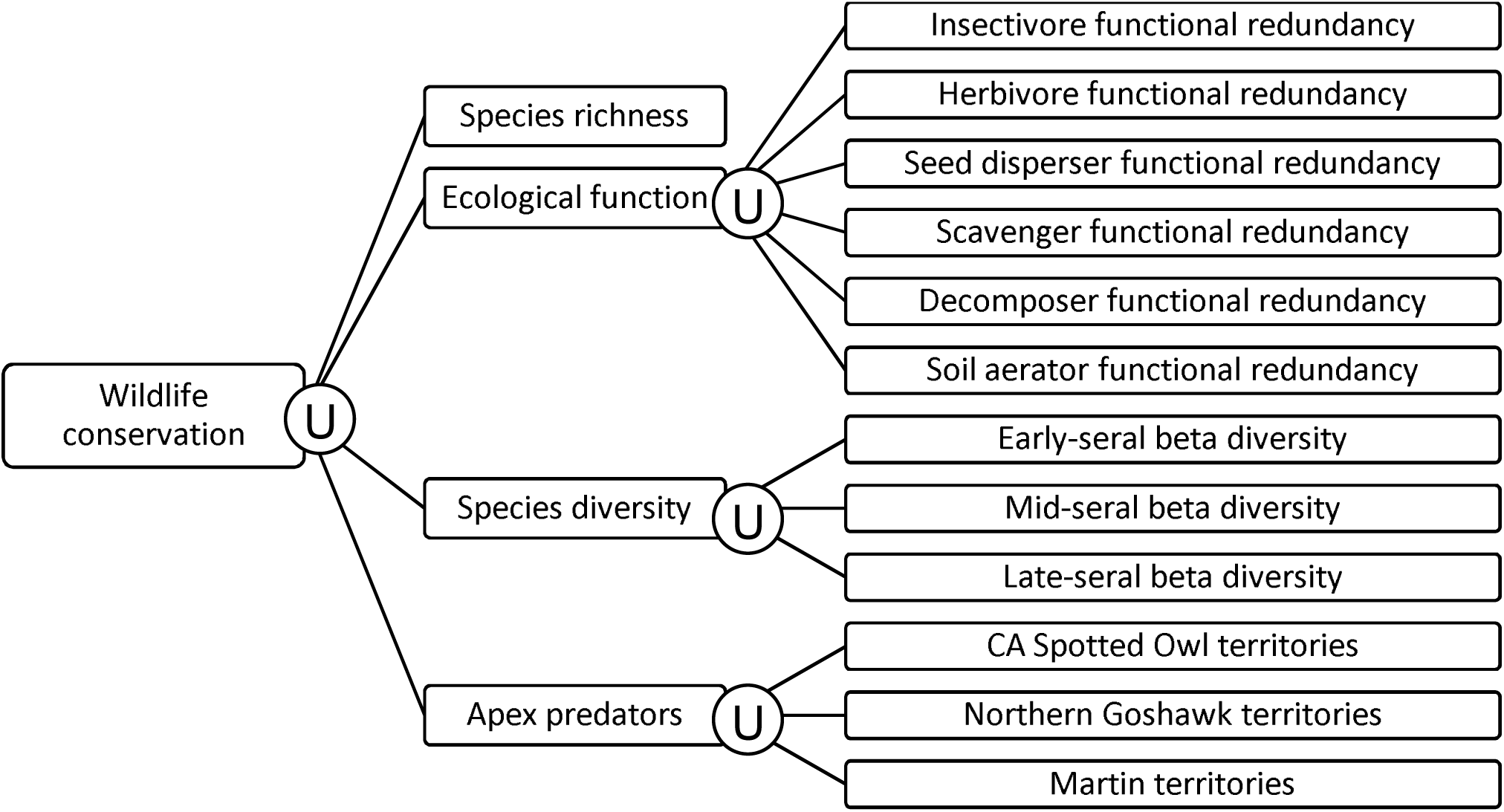
Logic model - wildlife conservation topic. Operators are depicted as either a “U” or an “A” in a circle to signify that a “union” or an “and” operator, respectively, was used.

##### 2.6.1.2. Quality water

Quality water has two components (Figure 3), phosphorous load and fine sediment load that employ a UNION operator. Phosphorous load evaluates the proportion of the annual phosphorous load to a baseline phosphorus load resulting from a landscape with no disturbance. Fine sediment is evaluated using the annual fine sediment under 16 microns relative to fine sediment under undisturbed conditions.

**Figure 3.**
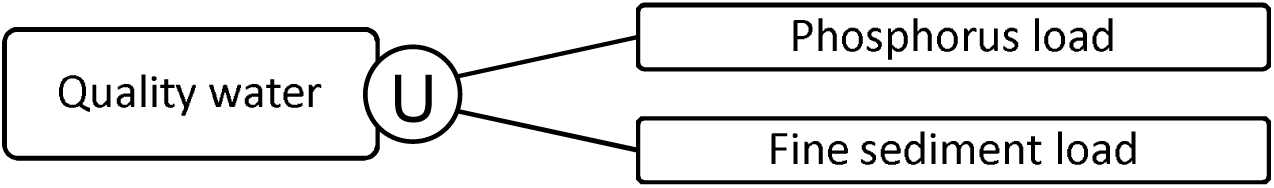
Logic model - quality water topic. Operators are depicted as either a “U” or an “A” in a circle to signify that a “union” or an “and” operator, respectively, was used.

##### 2.6.1.3. Upland vegetation health

Upland vegetation health is comprised of 13 data inputs in four main subtopics (Figure 4), and exclusively uses UNION operators. The Big trees subtopic evaluates the number of hectares with one or more trees over 150 years old. Forest cover evaluates the percent of vegetated area that is dominated by conifer, hardwood, and shrub. Seral stage is split into high elevation and mid elevation; both elevational levels evaluate the landscape percent that is in an early-, mid-, and late-seral stage. Finally, Composition evaluates the percent that yellow pine, white pine, and aspen represent of total biomass.

**Figure 4.**
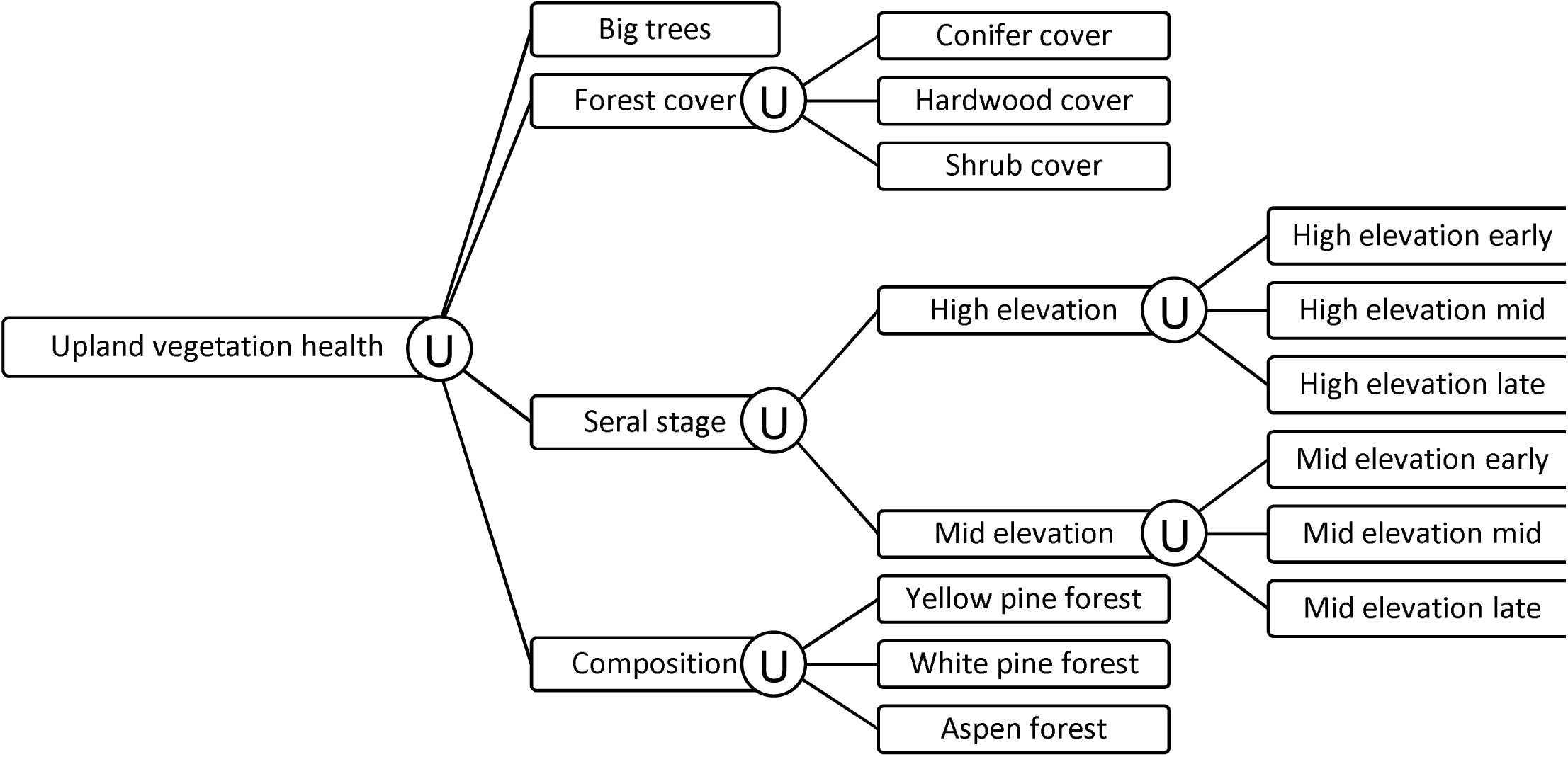
Logic model - upland vegetation health topic. Operators are depicted as either a “U” or an “A” in a circle to signify that a “union” or an “and” operator, respectively, was used.

##### 2.6.1.4. Functional fire

Functional fire uses UNION operators and is comprised of two subtopics (Figure 5). High severity patches evaluates the percent of landscape burned in high severity patches over 16 ha. Percent landscape burned evaluates the percent of the landscape that burns in low, moderate, and high severity per decade.

**Figure 5.**
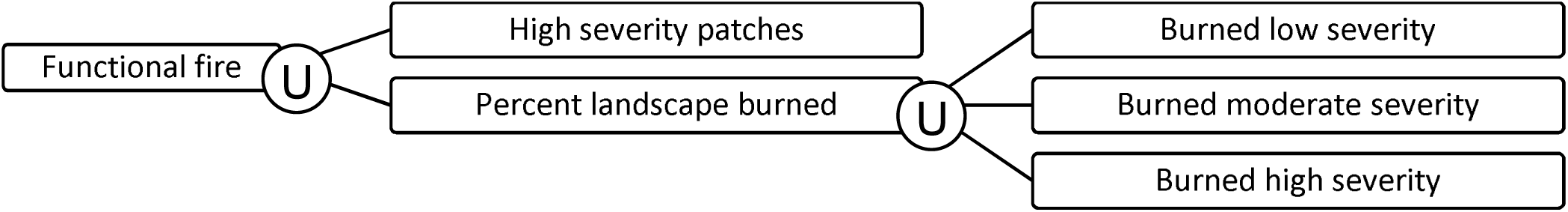
Logic model - functional fire topic. Operators are depicted as either a “U” or an “A” in a circle to signify that a “union” or an “and” operator, respectively, was used.

### 2.7. Multi-criteria decision model

A multi-criteria decision model assessed the five management scenarios (using both input from the logic models (Figures 2 to 5), as well as data for water quantity that came directly from topic area experts, and provided a score for each management scenario, by decade, to reflect how well each scenario performed with respect to achieving environmental quality and ecological resilience over the 100-year study period. The decision modeling step in the overall LTW analysis process is important in two respects. First, logic modeling was used to infer the state of forest landscape resilience for topic areas with many data dependencies. Second, the decision modeling component provides explicit modeling support for weighting topics that reflect the perspective of decision makers on the relative importance of decision criteria (Murphy, 2014). While the unweighted logic models assembled and assessed large quantities of complex data, the MCDM assessed the topics processed by the logic models in conjunction with topics that were generated by external science teams – weighted MCDM results were therefore a holistic view of the role that management scenarios had on modeled forest dynamics.

We used the MCDM component of EMDS, Criterium DecisionPlus (CDP) (*Criterium Decision Plus*, 2020), to evaluate the performance of the five alternative management scenarios (Murphy, 2014) with respect to maintaining or improving environmental quality. For large, complex, and abstract decision problems, as in the LTW application of EMDS, MCDMs are well suited to integrate both raw data and outputs from logic model evaluations (Figures 2 to 5). CDP ultimately outputs a score, between zero and one, that represents how well each scenario performs at each time point.

Decision models in CDP are goal oriented and implement the CDP analytic hierarchy process component (AHP) (Saaty, 1994, 1991) to choose among alternatives. In the current context, the goal is to select the best strategy for promoting environmental quality and promoting ecological resilience in the LTW study area. With implementation of CDP models in EMDS, lowest level criteria (also known as attributes) are evaluated by utility functions that map observed data values into a measure of utility with respect to satisfying the goal (Kamenetzky, 1982). Not all decision criteria are weighted equally by decision makers and EMDS permits criteria weighting to reflect real world perspectives of decision makers and managers with respect to their perceptions of the relative importance of the criteria and subcriteria. For example, in this ecologically driven assessment, the subject matter experts, and stakeholders that participated in designing the model, weighted wildlife conservation and water quality more heavily than upland vegetation health. While weights on decision model criteria may be assigned directly by managers, we used Saaty’s pairwise comparison methods and iteratively synthesized input from 24 project stakeholders (including agency managers, community stakeholders, and scientists) (Saaty, 1994, 1991).

#### 2.7.1. Multi-criteria decision model, model structure and data input

The decision model evaluates five ecosystem attributes that serve as the foundation for the CDP assessment (Table 1), and each enters into the CDP model as a unique data input. The latter table also includes information on whether a data input was pre-processed in the LTW EMDS logic models or provided as a direct input to the model from topic-area experts. Criteria and sub-criteria that make up the decision model are assessed in a three-level decision hierarchy (Figure 6).

**Table 1.**
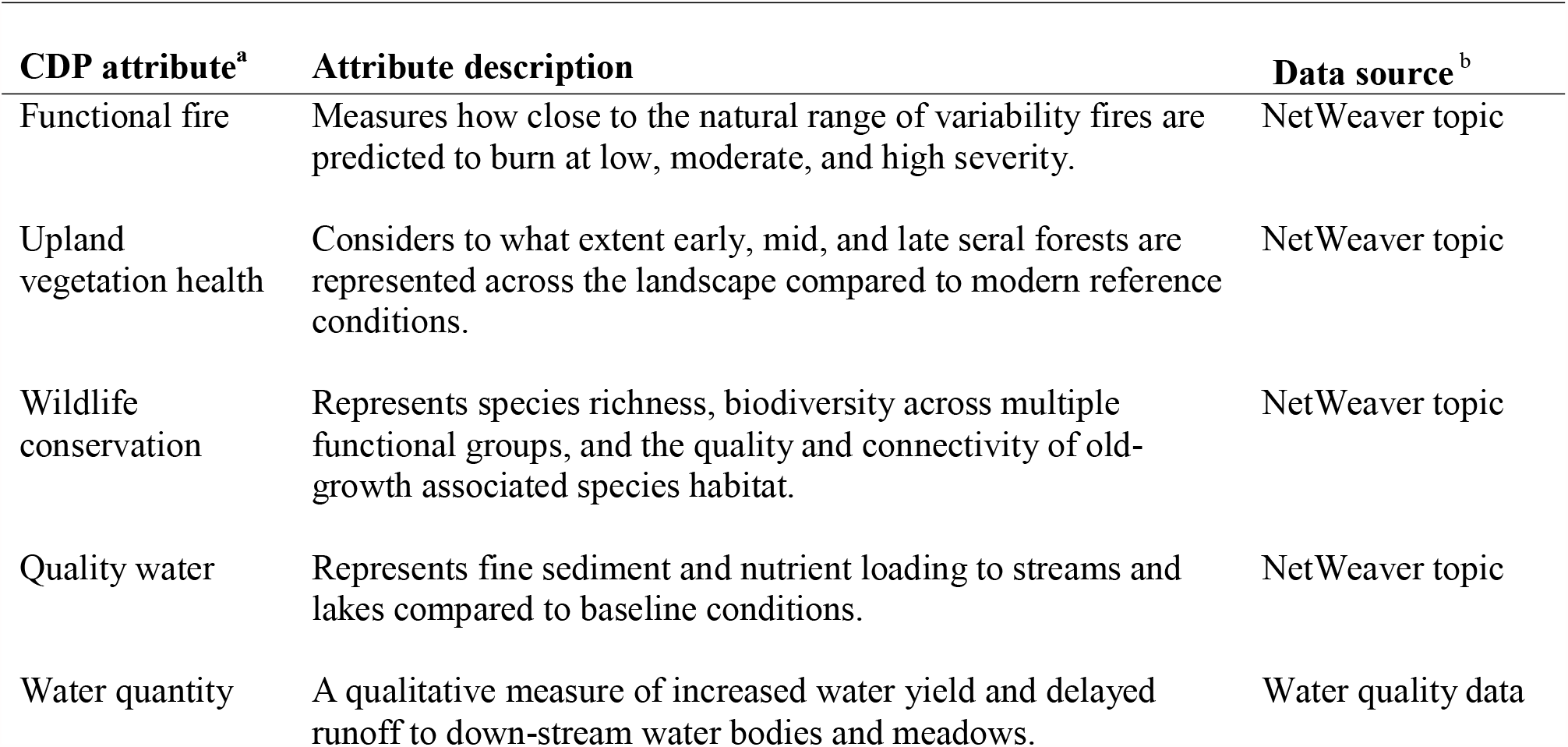
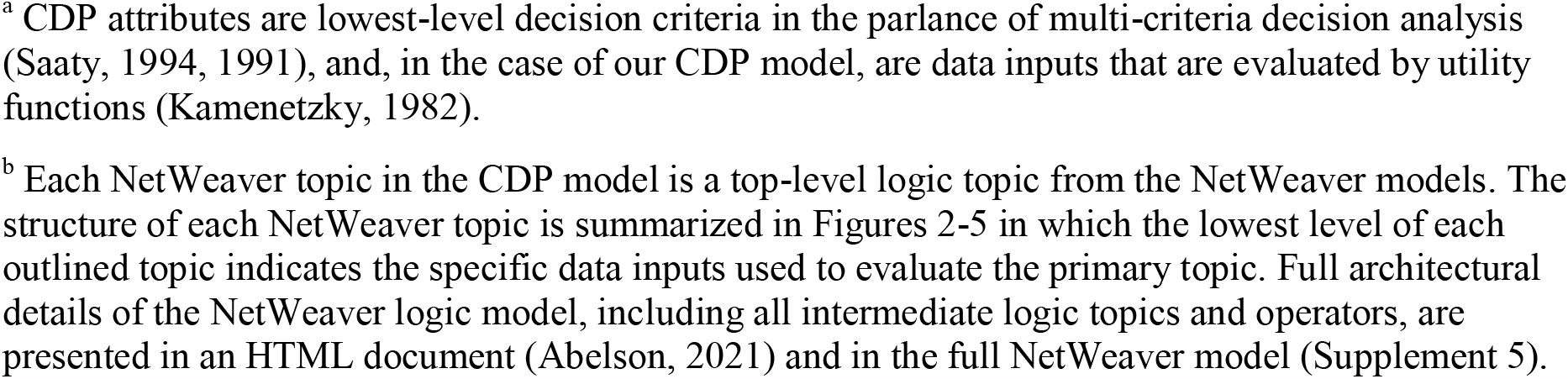
EMDS decision model topics and description

| **CDP attribute<sup>a</sup>** | **Attribute description** | **Data source<sup>b</sup>** |
| --- | --- | --- |
| Functional fire | Measures how close to the natural range of variability fires are predicted to burn at low, moderate, and high severity. | NetWeaver topic |
| Upland vegetation health | Considers to what extent early, mid, and late seral forests are represented across the landscape compared to modern reference conditions. | NetWeaver topic |
| Wildlife conservation | Represents species richness, biodiversity across multiple functional groups, and the quality and connectivity of old-growth associated species habitat. | NetWeaver topic |
| Quality water | Represents fine sediment and nutrient loading to streams and lakes compared to baseline conditions. | NetWeaver topic |
| Water quantity | A qualitative measure of increased water yield and delayed runoff to down-stream water bodies and meadows. | Water quality data |

<sup>a</sup> CDP attributes are lowest-level decision criteria in the parlance of multi-criteria decision analysis (Saaty, 1994, 1991), and, in the case of our CDP model, are data inputs that are evaluated by utility functions (Kamenetzky, 1982).
<sup>b</sup> Each NetWeaver topic in the CDP model is a top-level logic topic from the NetWeaver models. The structure of each NetWeaver topic is summarized in Figures 2-5 in which the lowest level of each outlined topic indicates the specific data inputs used to evaluate the primary topic. Full architectural details of the NetWeaver logic model, including all intermediate logic topics and operators, are presented in an HTML document (Abelson, 2021) and in the full NetWeaver model (Supplement 5).

**Figure 6.**
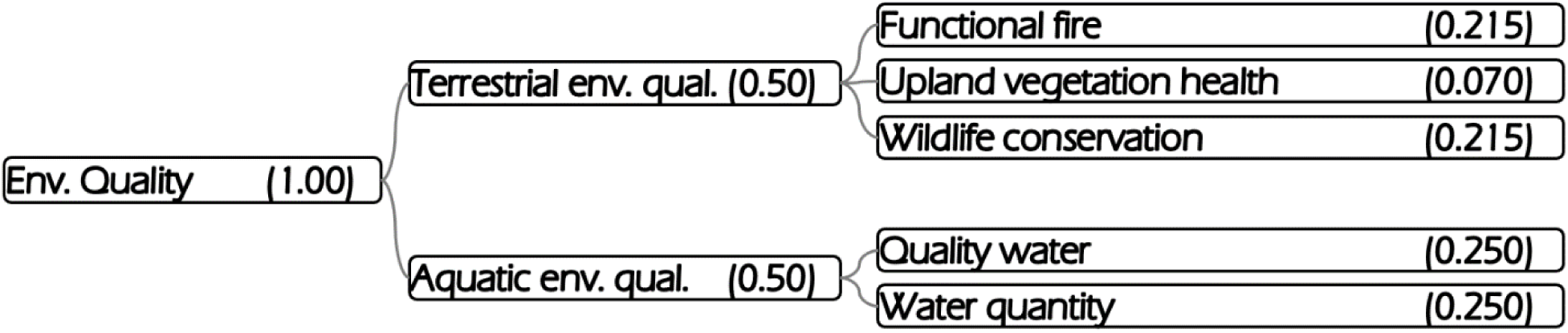
Decision model and hierarchy. CDP weights follow topic in paratheses; each column sums to 1.

To assess each management scenario, the LTW decision model evaluated two primary criteria that were weighted equally (Figure 6): terrestrial environmental quality and aquatic environmental quality. Terrestrial environmental quality broadly considers fire (Figure 5), vegetation (Figure 4), and wildlife (Figure 2). Aquatic environmental quality considers quality water and water quantity (Figure 3).

There may be several to many criteria worthy of consideration in the mind of a decision maker when examining any given decision, but not all criteria are necessarily equally important when compared to each other, and different contributors to model design are likely to bring different perspectives. In the next paragraphs we discuss 1) the composition of the team that participated in decision model design, 2) the method used to derive criterion weights, and 3) the iterative process used to arrive at a consensus on weights.

To derive criterion weights for the decision model, a group of 24 people were convened to anonymously provide their perspectives on the relative importance of the criteria. These 24 individuals were drawn from four LTW advisory groups: the interagency design team (agency-based technical experts), the stakeholder community committee (representatives of a diversity of interests in the basin), the stakeholder science committee (subset of the stakeholder community committee), and the environmental review team (agency staff charged with developing environmental documents to support management decisions). Of the 24, 12 were on the interagency design team, four were on the stakeholder community committee, six were on the stakeholder science committee, one was on both the interagency design team and the environmental review team, and one was on both the stakeholder community and stakeholder science committees.

CDP offers two basic methods to provide user input on weights for decision criteria: decision makers can directly assign weights to criteria, or Saaty’s pairwise comparison process (Saaty, 1994, 1991) can be used to derive the weights. In the AHP literature, the pairwise process is most typically preferred because it provides a systematic way to reason about the relative importance of criteria, and it provides a consistency metric that gives valuable feedback to participants on the consistency of their collective importance judgments. For design of the LTW decision model, we used a variant of the original Saaty pairwise methods called abbreviated pairwise. An impartial moderator facilitated meetings with participants to discuss and assign final consensus comparisons for each of the 15 possible pairwise comparisons needed for the abbreviated pairwise version.

The CDP score for each alternative management scenario in our CDP model is a measure of the utility of that scenario for satisfying the performance criteria for environmental quality. Model specifications for the utility functions and AHP are described in detail in Supplement 1 with values and model available in Supplement 6. Given that utility is measured on a scale of [0, 1], the utility score ranges of [0, 0.2], (0.2, 0.4], (0.4, 0.6], (0.6, 0.8], and (0.8, 1.0] can be interpreted as very low, low, moderate, high, and very high utility, respectively. Furthermore, because the utility score is addressing utility for satisfying model requirements of environmental quality, the same ranges can be interpreted as evidence of the degree to which modeled on-the-ground conditions are meeting resilience conditions (very low to very high resilience, Table 2).

**Table 2.** Interpreting MCDM performance scores.^a^

| Decision model performance score | Description |
| --- | --- |
| 0.8 – 1.0 | Excellent forest conditions |
| 0.6 – 0.8 | Good forest conditions |
| 0.4 – 0.6 | Intermediate forest conditions |
| 0.2 – 0.4 | Poor forest conditions |
| 0.0 – 0.2 | Very poor forest conditions |

<sup>a</sup> While performance scores for any given scenario (at any given time point) can be considered relatively against other scenarios at that same time point, these values can also be interpreted in absolute terms.

### 2.8. Output summary and analyses

The EMDS workflow evaluated 1,650 data inputs and resulted in 50 assessments of environmental quality that were plotted for each of the five management scenarios (Section 2.4) over ten time steps. We calculated summary statistics (i.e., min, max, mean, and standard deviation) of performance scores for each scenario over the ten decadal time points. We assemble these metrics for environmental resilience as well as for the two sub-topics that make up environmental quality: terrestrial and aquatic environmental quality (Figure 6).

## 3. Results

We evaluated 1,650 data inputs to calculate the outcomes for environmental quality resulting from five alternative management scenarios, each starting from the same ecological conditions, that were dynamically modeled over a century (and assessed at ten decadal time points). Each management scenario received a MCDM performance score for each time point. In addition to using performance scores to consider relative forest conditions, MCDM performance scores can also be interpreted in absolute terms (Table 2) where values closest to one are optimal as defined by the model parameters. All MCDM performance scores are included in Supplement 7.

Performance values close to one reflect optimal forest conditions and one can broadly group values from zero to one using breakpoints each 0.2 units.

We found that management activities are predicted to have important impacts on environmental quality over the 100-year time span (Figure 7). In relative terms, Scenario 5 outperformed all other scenarios and, in absolute terms, the mean performance score across the century for Scenario 5 was 0.87 (SD = 0.03) indicating that Scenario 5’s management activities promoted excellent outcomes with respect to environmental quality. Management activities prescribed by Scenarios 1 through 4 all result in good outcomes averaged over the century, resulting in good overall outcomes under these scenarios (MCDM performance scores between 0.6 and 0.8).

**Figure 7.**
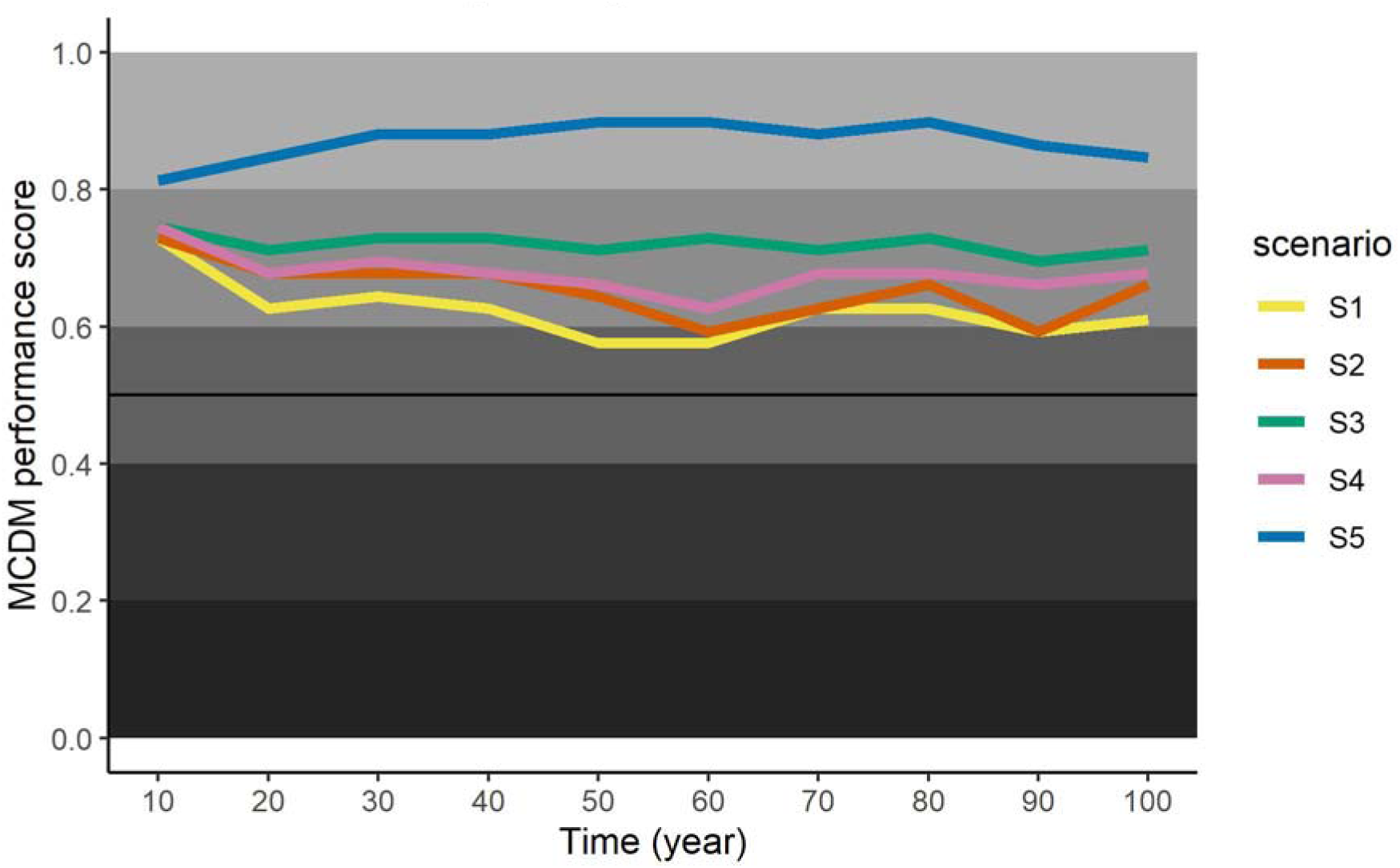
Multi-critera decision model results – Environmental quality.

Inter-year variability of forest condition is the degree to which outcomes change from decade to decade. While absolute outcomes are important, management scenarios outcomes that are robust to periodic events is often desirable. We found that Scenarios 1 and 2 were predicted to result in environmental quality outcomes with the most inter-year variability (standard deviation of 0.04 for both scenarios), while Scenario 3 had the least (standard deviation of 0.01). Scenarios 4 and 5 had similar inter-year variability in outcomes (standard deviation of 0.03 for both scenarios). In absolute terms, Scenarios 1 and 2 had similar minimum (0.58 and 0.59 respectively) and maximum values (0.73 for both scenarios) that were substantially below the Scenario 5 minimum of 0.81 to a maximum of 0.90.

Another important consideration is the comparison of future environmental quality and ecological resilience based on current management activities and alternative scenarios. Scenario 2 most closely resembles current management practices in the study area. We found that Scenario 2 is modeled to result in both variable and good outcomes over the coming century, though conditions dip into intermediate conditions for two of the 10 decadal periods considered here. We found that alternative management approaches, especially Scenarios 5 (mean performance score of 0.87) and 3 (mean performance score of 0.72), resulted in considerably better environmental quality than Scenario 2 (mean performance score of 0.65), with reduced variability in inter-year outcomes (standard deviation in performance score of Scenario 5 and 3, respectively, are 0.03 and 0.01, while standard deviation of performance scores for Scenario 2 was 0.04).

Scenario performance is equally dependent upon terrestrial and aquatic environmental quality (Figure 6). It is possible to better understand scenario performance values by examining themes in the MCDM scenario performance values for terrestrial (Figure 8) and aquatic environmental quality (Figure 9) across the 100-year period for which forest conditions were modeled. The minimum MCDM performance scores for terrestrial environmental quality, across all scenarios and time points, was 0.7 compared to a minimum aquatic environmental quality score of 0.4. All scenarios performed well in terms of terrestrial environmental quality and this compensated for the declining mid- to late-century performance of scenarios one, two, and four in aquatic environmental quality. In terms of terrestrial environmental quality, Scenario 5 was the leading scenario with a mean MCDM scenario performance score of 0.9 (SD 0.04) followed by Scenario 3, which also is modeled to have minimal inter-year variability (mean = 0.81, SD 0.02) and Scenario 4 with a similar mean performance score but with increased inter-year variability (mean = 0.8, SD = 0.03).

**Figure 8.**
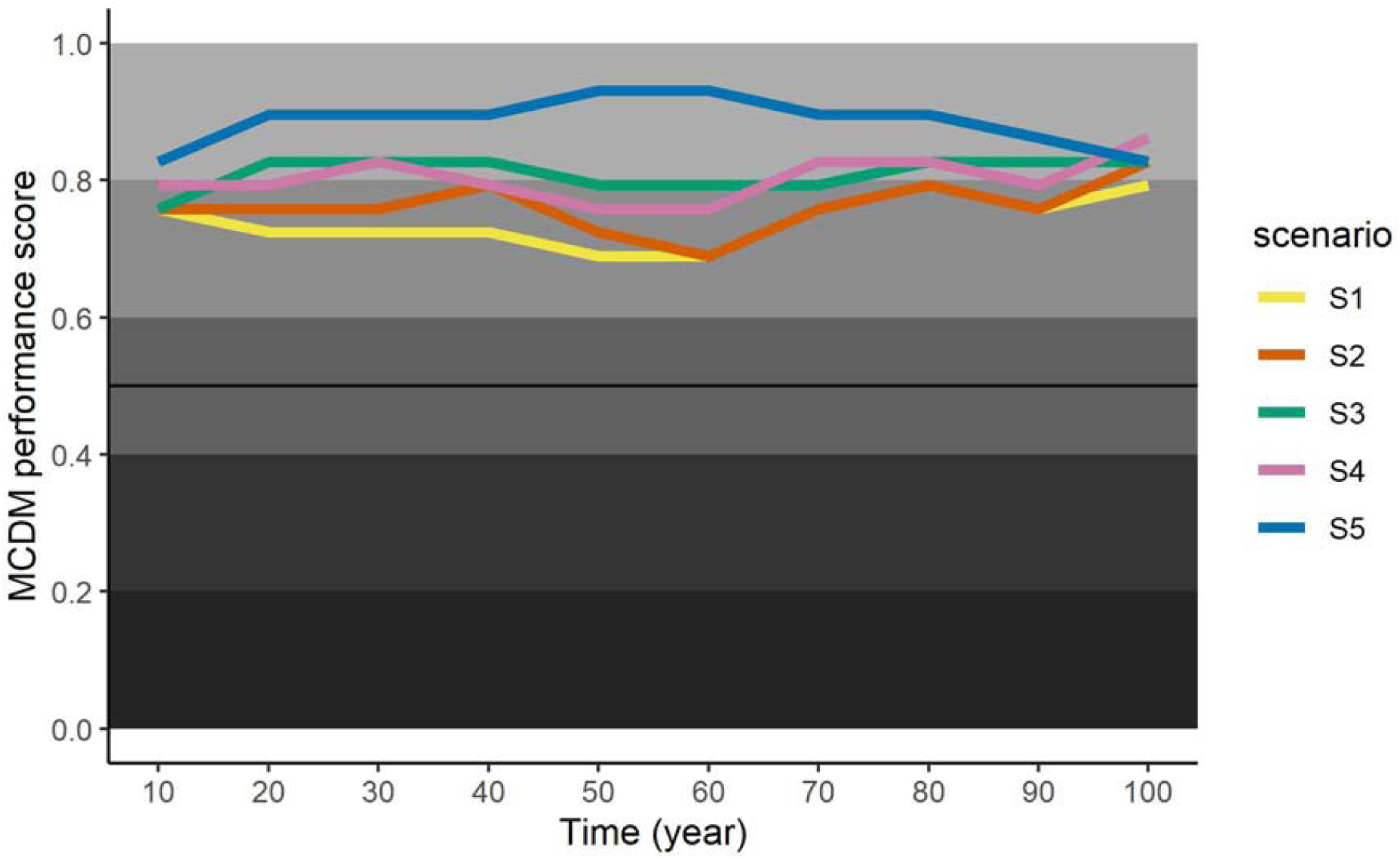
Multi-criteria decision model results – Terrestrial environmental quality.

**Figure 9.**
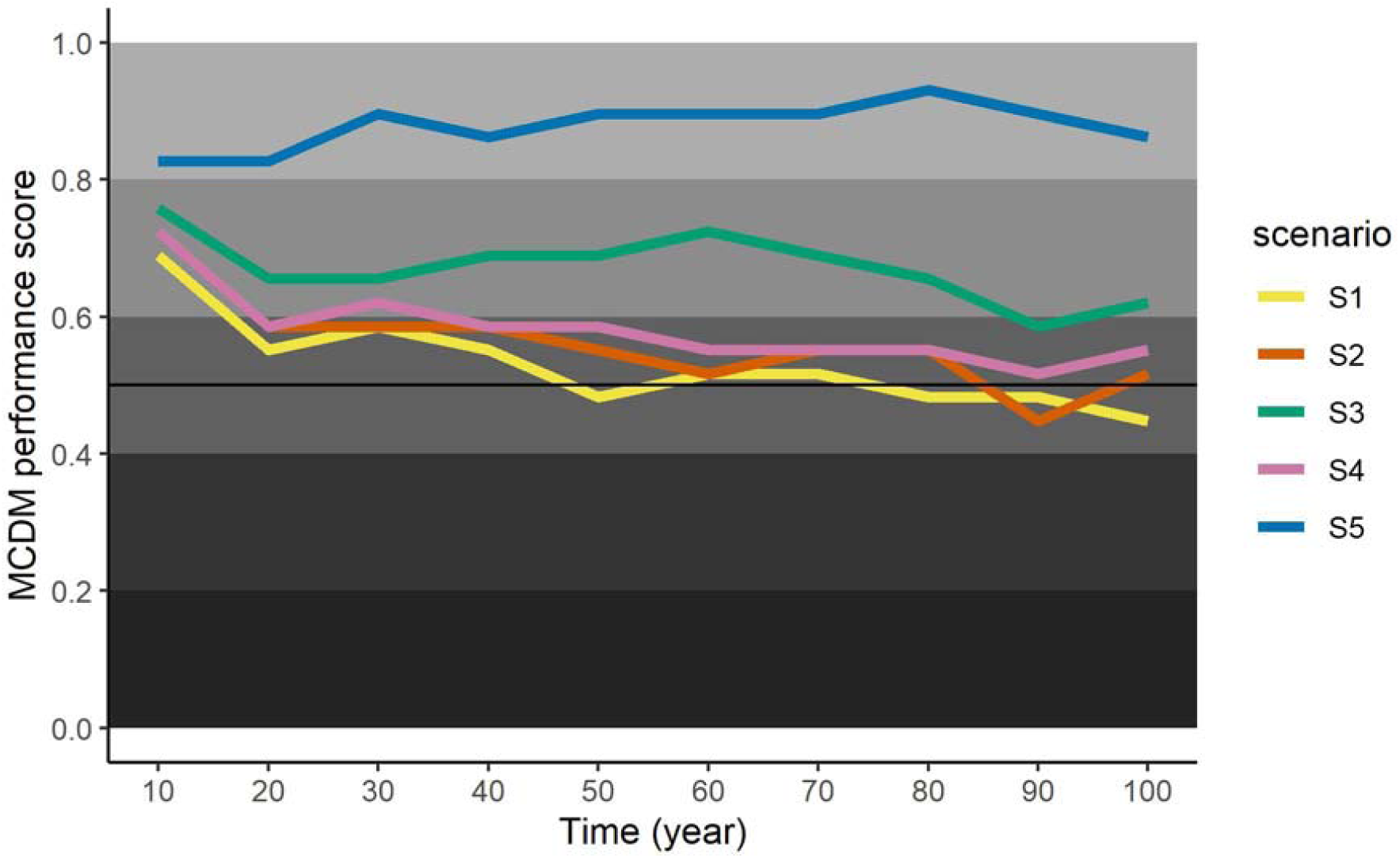
Multi-criteria decision model results – Aquatic environmental quality.

Differentiation between management scenarios was more pronounced in aquatic environmental quality (Figure 9) than terrestrial environmental quality (Figure 8) when looking at trajectories of performance of the scenarios over time. Considering only the aquatic environmental quality criterion, Scenario 5 was predicted to result in excellent environmental quality outcomes (i.e., those with MCDM performance scores between 0.8-1.0), Scenario 3 resulted in generally good outcomes (i.e., performance scores between 0.6-0.8), while Scenarios 1, 2, and 4 are expected to result in generally intermediate performance scores (i.e., scores between 0.4-0.6).

## 4. Discussion

EMDS, for the Lake Tahoe West project, provides guidance to decision-makers on the performance of five management scenarios over a century on modeled landscape conditions. We find first that management approach influences predicted landscape condition over time. Since management approach results in varied landscape conditions, decision-makers are faced with the mandate to consider five management scenarios, and the forest conditions that result from each of 1,650 individual data inputs. This challenge is too extensive for any person to sensibly consider, underscoring the need for a tool to process these data in a way that mimics the approach a decision-maker would take. EMDS results represent the forest conditions, over time, for each of the five management scenarios.

We propose three main evaluation considerations when considering LTW EMDS output. Because these results span 100 years, we propose that it is important to evaluate 1) the relative performance of one scenario versus another at any given time point, 2) absolute scenario performance at each time point, 3) year-to-year variability and maximum/minimum performance, and 4) predicted forest condition based on current practices compared to alternatives. Relative performance elucidates the trade-offs between choosing one scenario over another. Absolute scenario performance identifies the forest conditions at any given time point and for any given scenario. Assessing absolute performance is important because while one scenario may outperform all the others, it still may result in “unacceptable” landscape conditions. Finally, year-to-year variability could be undesirable in and of itself or may result in dips below an “acceptable” forest condition threshold.

Regarding the four evaluation considerations, we find that intensive fire-based management (Scenario 5) outperforms the other four scenarios by generally cultivating landscapes in excellent condition with minimal between-year variation. Management approaches utilizing forest thinning in and outside of the WUI, using biomechanical (Scenario 3) removal techniques is the next best option in terms of absolute performance, relative performance, and between year variability. In absolute terms, the other management scenarios that range from only suppressing naturally occurring fires to management calling for moderate levels of biomass removal result in good forest conditions. Those scenarios with the least amount of management (i.e., Scenarios 1 and 2), including the management scenario that most closely resembles current management activities (Scenario 2), comparatively performed the most poorly with decades of intermediate forest conditions and higher inter-year variability then the other management options.

The utility of EMDS in the Lake Tahoe West project is not limited to numerical valuations of forest conditions over time. An important contribution of EMDS is to the *process* of how forest conditions are evaluated when working in a context of multiple science areas and stakeholders with divergent approaches to forest management. The EMDS process facilitated four main conversation topics: 1) which topic areas should be included, 2) how topic areas should be structured into a topic hierarchy (topic hierarchy directly influences evaluation), 3) topic area data evaluation (for example, a determination of the “ideal” amount of rainfall – defining the ideal and suboptimal inches of annual rainfall), 4) topic area weighting. Each of these conversation topics required many meetings with different stakeholders, were iterative (i.e. a paradigm to move forward was arrived at by constructing a proposal, reviewing with stakeholders, revising, and then repeating the previous three steps).

The *process* of the EMDS LTW project is itself an important result. The EMDS process provided structure to a large, unstructured, and intractable challenge. EMDS established data, interpretation, and analysis needs that served as the backbone for conversations among experts in different fields along with stakeholders. The process of this project is itself a result as the EMDS is not pre-ordained and instead required effort and discussion among a diverse group to provide necessary base information to parameterize the EMDS model. The EMDS process for LTW is embodied in the logic models (Figures 2-5), the decision model hierarchy (Figure 6), FMFs of the logic models (Supplement 5) and decision model utility functions (Supplement 6).

We demonstrate here that EMDS is a decision support tool that is well suited to an aspatial ecological analysis of complex, varied, and multi-dimensional data. Additionally, we also present a use case for new components of EMDS (i.e. Wexflow) that facilitated our analyses. The Wexflow workflow editor simplified a big project (i.e. 60 individual EMDS model runs) into a tractable series of routines (i.e. a single “unified” model). EMDS is traditionally used to evaluate spatial data; here we demonstrate that it is also an equally valuable tool when spatial data is distilled into aspatial data. Finally, we demonstrate here that EMDS, as a decision support tool, is equally capable at handling data across long time-horizons as it is handling a landscape at a single point in time.

## 5. Endnotes

The use of trade or firm names in this publication is for reader information and does not imply endorsement by the U.S. Department of Agriculture of any product or service.

### interests

The authors declare no competing interests.

## Acknowledgments

Amelia Wolf, Angela White, Forest Schafer, Sarah Di Vittorio, Dorian Fougeres, Jonathan Long, Brian Garrett, Shana Gross, Jen Greenburg, Forest Schaffer, Nadia Tase, and Jason Vasquez.

**Supplement 1. EMDS, NetWeaver, Criterium DecisionPlus, and Wexflow Methodological background and details**.

**Supplement 2. EMDS workflow automation for LTW project implemented using Wexflow.**

**Supplement 3. LTW decision support tool raw input data**.

**Supplement 4. NetWeaver logic model;**. **nw2 file format version for use in NetWeaver software**

**Supplement 5. NetWeaver model fuzzy membership function values. These values are used for interpreting raw input data in the NetWeaver model and serve as input data for the NetWeaver model in Supplement 4**.

**Supplement 6. Criterium DecisionPlus model;**. **cdpx file format version for us in Criterium DecisionPlus**.

**Supplement 7. Performance scores for environmental quality, terrestrial environmental quality, and aquatic environmental quality (i.e**., **Criterium DecisionPlus output)**.

