## Supplemental info 1 for "Strategic decision support for long-term conservation management planning"

**Supplement 1. EMDS, NetWeaver, Criterium DecisionPlus, and Wexflow Methodological background and details.**

**EMDS background**

The Ecosystem Management Decision Support (EMDS) system is a state-of-the-art modeling framework for spatial decision support of environmental analysis and planning at multiple geographic scales (Reynolds and Hessburg, 2014). The system integrates modern geographic information system (GIS) as well as logic-based reasoning and a variety of decision modeling technologies to provide explicit, practical decision support for strategic and tactical planning. In the LTW project, data inputs were evaluated in EMDS to assess the degree to which each of five management scenarios were expected to provide optimal conditions for forest landscape resilience.

Because EMDS is a generic solution framework, it can be applied to an extremely broad array of problems at virtually any and all spatial scales. Examples of the breadth of decision support solutions provided by EMDS can be found on the EMDS Wikipedia page (“Ecosystem Management Decision Support,” 2020).

While EMDS is designed as a spatial decision support tool version 8.0 is implemented as a spatial decision support plug-in component to both ArcMap^[1]^ (ArcGIS Desktop, 2020) and the open-source GIS, QGIS (QGIS Geographic Information System, 2020)

**Logic model**

A logic model can be thought of as a type of meta-database that is ontologically committed to a logic-processing engine (Guarino, 1998). In the LTW project, as in most natural resource management projects with a high level of public interest, black box solutions are a political liability. Ultimately the logic processor displays evaluated states of complex logic models in a highly intuitive interface which we used as a powerful communication tool to explain results to broad audiences.

The logic model assesses data in two main ways. First, the raw data (per time point and management scenario) were read and logically evaluated with fuzzy membership functions (FMFs) (Saunders and Miller, 2014) that translate observed input values into a measure of the strength of evidence that the observed value (in the case of LTW logic model) contributes to forest landscape resilience (i.e., the -1 to 1 range described above). Subsequently, we refer to logic topics that process inputs with FMFs as elementary topics. For the LTW logic model, each FMF was specified as a simple ramp function defined by two x-y pairs, x_1_y_1_and x_2_y_2_, in which x_1_ specifies the observed value at which there is no evidence for resilience (y_1_=-1) and x_2_ specifies the observed value at which there is full evidence for resilience (y_2_=1). LTW project scientists, who were responsible for each of the primary logic topics, established the x_1_and x_2_ values for the FMFs covered under their primary logic topic. Technically, each FMF is a compound function consisting of three parts:

y=-1 when x≤x_1_,

y=1 when x≥x_2_, and

y is linearly interpolated in the open interval (x_1_, x_2_).

Given the NetWeaver resilience scores as calculated by the elementary topics, NetWeaver propagates evidence scores upward through successive levels of the logic structure (logic structures for LTW topics space are described in the following section) with logic operators that synthesize outputs from child (i.e., lower level) topics. NetWeaver includes a variety of operators for logical synthesis, but, in the case of the LTW NetWeaver logic model, all syntheses were performed with either the AND or UNION operators. The UNION operator averages evidence scores from two or more lower-level topics, while the AND operator uses a minimum-biased weighted average. The UNION operator is appropriate when its child topics have a compensatory effect on the evaluation of the parent topic; in other words, negative evidence can be compensated by positive evidence The AND operator is appropriate when its child topics are considered limiting factors as opposed to compensatory. The effect of the minimum-biased calculation is to heavily weight the evidence toward the most negative line of evidence, although any line of evidence having a value of -1 is a special case, in which case the result of the AND operator is also -1.

**Multi-criteria decision model, implemented in CDP**

Given the apparent similarity in the functionality of logic and decision models, some readers may wonder about the rationale for the use of both these tools in the EMDS framework. Earlier, we alluded to part of the answer: the two types of models can be used in complementary fashion to tease apart issues of system state versus priorities for management action. In the process of doing so, this makes it possible to develop two smaller and simpler models. In effect, we used logic models to distill copious detailed ecological information, and avoided designing very large decision models, which in our experience can become cumbersome and impractical. Perhaps at least as importantly though, logic models can easily model large, complex problems in which at least some of the dependencies modeled may be nonlinear, and thus difficult to handle in a decision model, which is intrinsically linear.

For example, in the “social values” topic the abbreviated pairwise method requires three comparisons (i.e., there are four sub-topics in “social values” – the abbreviated pairwise method would ask topic A vs. B, A vs. C, and C vs. D) rather than six (i.e., topic A vs. B, A vs. C, A vs. D, B vs. C, B vs. D, and C vs. D). A side effect of the abbreviated pairwise method is that the CDP software computes the remaining (missing) comparisons so that the consistency metric is perfect or at least nearly so.

Whereas the full pairwise method requires making all possible pairwise comparisons among subcriteria of a criterion, the abbreviated pairwise method selects a minimal spanning set of pairwise comparisons that can greatly reduce the number of comparisons that need to be provided.

At the start of the rating process, each participant individually scored each pairwise comparison among subcriteria at a given level of the model hierarchy using a modified version of the AHP comparison scale of 1 to 9, in which a value of 1 means that criteria A and B are equally important, and a value of 9 means that criterion A is absolutely more important than B. To simplify choices for respondents (comparing A to B), we used the odd numbers on the scale to approximate a Likert scale as follows: 1,3, 5, 7 and 9 mean equally important, weakly more important, definitely more important, strongly more important, and absolutely more important, respectively. We also allowed for an “I don’t know” response, in which case the observation was omitted from subsequent tallies and reporting.

After each participant had an opportunity to review and provide their input on each pairwise comparison, we then assembled, summarized, and generated histograms of all responses (i.e., how many respondents indicated that topic A is absolutely more important than topic B, etc.).

Resulting values were entered directly into CDP to complete computation of the final criterion weights (Figure 10). From the latter figure, note that the weights on the 16 attributes (lowest-level criteria) sum to 1. The final decision score for a scenario for each decadal period is calculated in CDP as the sum of the products of an attribute’s criterion weight and its utility value.

**Wexflow workflow editor**

The third EMDS component that we employ in our LTW application is a customized version of the open source Wexflow workflow editor (El Assas, 2020), which was introduced in EMDS version 8 and was used to automate analyses.

The LTW application is the first published EMDS example of an application that illustrates how workflows can automate a complex analytical process. Briefly, the overall LTW analysis required running a NetWeaver logic model 50 times (10 time steps by five alternative scenarios) and then using those NetWeaver outputs iteratively in 10 distinct CDP models to compare each of the five scenarios at each time step. Heretofore, that would have required an analyst to spend considerable time running the total of 60 model runs. However, using workflows designed in Wexflow, streamlined the entire set of analyses.

The concept of workflows is relatively straightforward. A workflow editor allows users to graphically assemble sequences of actions that have been individually programmed in a variety of languages, as sequences of discrete processing steps to be performed to accomplish some analytical task. A sequence of activities constitutes a workflow, and the workflow concept is extensible to include sequences of workflows (i.e,, workflows of workflows). Workflows are perhaps best known as a way to automate (geo)processing tasks, but our customized version of Wexflow extends the core automation feature of workflows to include a highly interactive workflow interface in which the editor can be programmed to interact with system users in real time to adjust parameters and process pathways at one or more intermediate steps in a complex workflow sequence. Complementing its ability to design and execute workflows, the Wexflow component in EMDS version 8 also interoperates with a workflow library manager that allows the storage, retrieval, and sharing of workflows across EMDS applications.

1. The use of trade or firm names in this publication is for reader information and does not imply endorsement by the U.S. Department of Agriculture of any product or service.
